## Supplemental Text S1 for "MICOM: metagenome-scale modeling to infer metabolic interactions in the gut microbiota"

September 18, 2019

### 1 Effect of regularization on growth rates

We consider a community model with growth rates $\mathbf{g} = \{g_1, \ldots, g_n\}, g_i \geq 0$ and relative abundances $\mathbf{a} = \{a_1, \ldots, a_n\}$. The maximal community growth rate is given by $\mu_c^* = \max \mathbf{a} \cdot \mathbf{g}$ again and we will apply a tradeoff parameter $\alpha \in (0, 1]$.

In the decoupled microbial community model species do not interact, share no metabolite imports and do not experience resource limitation. Consequently, all $g_i$ are independent of each other. During regularization we will minimize a regularization function $r(\mathbf{g}) \in \mathbb{R}^+$ with the previously described tradeoff condition:

$$\mu_c = \mathbf{a}^T \mathbf{g} \geq \alpha \mu_c^*$$

If the regularization function is strictly monotonic in $\mathbf{g}$ this inequality will be an equality. In general, finding an analytical solution for the fullk set of equality and inequality constraints is difficult and requires the evaluation of complementary slackness of the inequalities. However, on can find an optimal solution that only considers the optimality constraint and the regularization term and which describes the best case solution when all other constraints can be met.

This can be achieved with a Lagrangfe multiplier which induces the following minimization problem:

$$\min_{\mathbf{g},\lambda} L(\mathbf{g}, \lambda) = r(\mathbf{g}) - \lambda \left( \mathbf{a}^T \mathbf{g} - \alpha \mu_c^* \right)$$

In general we will only consider the non-trivial case where there exists at least one $i$ such that $g_i > 0$.

#### 1.1 Linear regularization

If $r$ is linear it can be expressed as $r(\mathbf{g}) = \boldsymbol{\beta}^T \mathbf{g} + c$. This induces the following Lagrangian optimization problem:

$$\min_{\mathbf{g},\lambda} L(\mathbf{g}, \lambda) = \boldsymbol{\beta}^T \mathbf{g} + c - \lambda \left( \mathbf{a}^T \mathbf{g} - \alpha \mu_c^* \right)$$

with the following conditions for the existence of a minimum:

$$\forall i : \frac{\partial L}{\partial g_i} = \beta_i - a_i\lambda = 0$$
$$\mathbf{a}^T\mathbf{g} - \alpha\mu_c^* = 0$$

Thus we have to fulfill:

$$\forall i : \lambda = \frac{\beta_i}{a_i}$$
$$\mathbf{a}^T\mathbf{g} = \alpha\mu_c^*$$

As we can see the first equation is independent of $g_i$ and only imposes conditions on the ratio between coefficients and abundance whereas the second condition can be fullfilled by many different sets of growth rates. So linear regularization either has no defined minimum or infinitely many minima. In particular it does not guarantee that $a_i > 0 \iff g_i > 0$.

#### 1.2 Quadratic regularization

We will now consider a simple quadratic regularization (L2) with $r(\mathbf{g}) = \mathbf{g}^T\mathbf{g}$. This induces the following Lagrangian optimization problem:

$$\min_{\mathbf{g},\lambda} L(\mathbf{g}, \lambda) = \mathbf{g}^T\mathbf{g} - \lambda\left(\mathbf{a}^T\mathbf{g} - \alpha\mu_c^*\right)$$

with the following conditions for the existence of a minimum:

$$\forall i : \frac{\partial L}{\partial g_i} = 2g_i - a_i\lambda = 0$$
$$\mathbf{a}^T\mathbf{g} - \alpha\mu_c^* = 0$$

Thus we have to fulfill:

$$\forall i : g_i = \frac{\lambda}{2}a_i$$
$$\mathbf{a}^T\mathbf{g} = \alpha\mu_c^*$$

Plugging the first into the second equation gives:

$$\frac{\lambda}{2}\mathbf{a}^T\mathbf{a} = \alpha\mu_c^*$$
$$\rightarrow \lambda = \frac{\alpha\mu_c^*}{\mathbf{a}^T\mathbf{a}}$$

Pluggin this into the previous equation for $g_i$ now yields:

$$\forall i : g_i = \frac{\alpha\mu_c^*}{\mathbf{a}^T\mathbf{a}}a_i$$

Thus, we have a well defined minimum, and since $\frac{\alpha\mu_c^*}{\mathbf{a}^T\mathbf{a}}$ is a constant, growth rates depend linearly on the abundances and it follows that $a_i > 0 \iff g_i > 0$.

#### 1.3 Conclusions

Thus, in the unconstrained case L2 regularization will yield growth rates that are perfectly correlated with abundances. However, based on additional constraints, especially resource limitation those optimal growth rates may not be reached which can result in different optima.
